# Allantoic fluid-based qPCR for early-onset *in ovo* sexing

**DOI:** 10.1101/2023.12.13.571516

**Authors:** Simão Santos, Matthias Corion, Bart De Ketelaere, Jeroen Lammertyn, Dragana Spasic

## Abstract

The culling of day-old male chickens remains an important welfare issue in the poultry industry. Several governments (*e.g.*, Germany, France or Italy) have prohibited this practice, pushing the hatcheries to look for alternatives. Although different solutions exist for solving this problem, sex determination during embryo’s incubation (so called *in ovo* sexing) is considered the most suitable both for consumers and the industry since it aligns with the increasing demand for ethical and sustainable agriculture. However, to be applied in the market, *in ovo* sexing technologies have to meet a number of requirements, such as being compatible with all egg colors and early developmental stages, while maintaining high hatchability rate and accuracy at low cost and high throughput. To meet these requirements, we studied the use of the sexual genes *HINTW* (female-specific) and *DMRT-1* (present in both males and females) between incubation days 6 and 9. By utilizing quantitative polymerase chain reaction (qPCR) as analysis method and allantoic fluid (AF) as sample, our study confirmed specific detection of these genes in AF, remarkably already at day 6 of development. In a blind study performed with 80 eggs, we achieved a 95% accuracy rate in sorting embryos when using the *HINTW* gene alone and an outstanding 100% accuracy rate when using Δλ values (difference between the *HINTW* and *DMRT-1* qPCR cycle threshold (Ct)). Importantly, the AF sampling procedure did not reveal any significant detrimental effects on hatchability or embryo development, revealing high potential for this understudied type of sample to be used. In conclusion, the developed assay can provide more in-depth information about AF as a sample for genomic *in ovo* sexing and open new industrial possibilities for developing faster and cheaper assays.

**Research highlights:**

- A highly reliable and accurate method for *in ovo* sexing in early incubation stages is established.
- Allantoic fluid can be easily extracted with minimal invasiveness while providing the necessary genomic material for sexual sorting.
- *HINTW* was found to be present in both female and male samples due to possible maternal contamination.
- *HINTW* alone or combined with *DMRT-1* gene enables early-onset sexing with 100% accuracy.

## 1. Introduction

Culling 1-day-old male chickens remains a standard practice within the poultry industry^1^. Approximately 372 million male day-old chicks are killed annually in the EU upon hatching^2^. This practice is attributed to the lack of purpose given to male chickens within the laying hen industry as they do not lay eggs and their meat is not appreciated ^3^. Within European countries, both consumers and industry are pressuring European governments to end the culling and to have food free of animal suffering. This is because 1) the consumers are increasingly aware of animal welfare and threaten to avoid poultry products^3^, while 2) the industry finds this practice unprofitable since there is no revenue from the male chicken embryos and the workers that handle the day-old chicken embryos dislike the gruesome practices^1^. Recently, Germany^4^, France^5^ and Italy^6^ have created legislations banning the culling practices and imposing sexual determination and egg separation before the potential embryo pain perception onset or growing the males, giving them the opportunity for a whole life^7^. Recent studies from Technische Universität München showed that pain perception is expected to start at day 14 of incubation due to the presence of encephalogram signals^8^.

Currently, there are three main strategies to avoid culling male chicken embryos: 1) genetically modified organisms (GMO), 2) dual-purpose chicken lines and 3) *in ovo* sexing. GMOs are currently not allowed in European products for human consumption^9^. Although previous work already showed that GMOs provide high sexing accuracies early in incubation^10^, for full implementation of GMO chickens (in countries outside the EU), the breeding companies will have to adopt a new pure chicken line (*i.e.*, with a tag in their genome). Moreover, besides the effects on the price, the animals’ and consumers’ long-term health and the industry reputation are still unknown^1^. Dual-purpose chicken lines present a higher financial and environmental burden since females and males have to be grown with more feed than the standard laying hen lines^11^. Thus, *in ovo* sexing (*i.e.*, sexual detection before hatching) is the practice that the industry prefers to resolve this issue since it avoids investing in male chickens and allows embryo disposal before pain perception development^3^.

However, for being applied in industrial settings, the technologies are required to meet several market requirements: 1) compatibility with all colors of eggs, 2) high throughput (> 20 000 eggs/hour, complying with the market’s high product demand), 3) high accuracy (> 98%), 4) applied early in incubation, imperatively before day 13 of incubation, 5) maintaining a high hatchability rate and 6) being low cost (around 2-3 €/day-old-chick; Bruijnis et al., 2015). Given the challenges presented by these requirements, significant efforts have been made by both industry and researchers to develop methods of *in ovo* sexing that can comply with all the above-mentioned requirements.

Current *in ovo* sexing technologies can be categorized as optical and non-optical^12^. On the one hand, optical techniques can use visible near-infrared spectroscopy to candle the egg on days 13 or 14, distinguishing the embryos’ feather colors, but these are only working on brown eggs with 99% accuracy^2^. This approach is currently the only commercial optical technique marketed by Agri Advanced Technologies (2020). Another optical technique is Raman spectroscopy, which, although it can be performed on day 3.5 with 96% accuracy, is highly invasive since it requires opening a 12.8 mm hole in the eggshell, decreasing the eggs’ hatchability rate^14^.

On the other hand, non-optical methods rely on invasive sampling (*e.g.*, blood, tissue, or allantoic fluid (AF)), followed by its analysis with 1) ELISA (for hormones)^15^, 2) PCR (for specific DNA sequences), or 3) mass spectrometry (*e.g.*, detection of valine and glucose). Although ELISA and PCR can be both considered minimally invasive when using AF as a sample, while offering high accuracy (*i.e.*, and 98 and 99.5%, respectively), they can be lengthy (with time-to-result of more than 60 minutes), thus decreasing the throughput and increasing the overall cost. Contrary to this, mass spectrometry enables short testing time (approximately 20 minutes), but low accuracies for industrial standards (90 – 95%). Despite the fact that no present sexing method can meet all the market requirements, several companies have been commercializing some of these techniques^16–18^. Nevertheless, because of the imposed legislations, there is a huge pressure in this field for new technologies that could fulfill all market demands.

Among non-optical techniques, DNA analysis is the most accurate one, allowing sex detection before day 8^19^, guaranteeing close to 100% accuracy. In birds, the sex chromosomes are represented by a ZZ chromosome pair in males and a ZW in females^20^, thereby allowing identification of female-specific sequences in the W chromosome. Previous reports have shown such detection using genes *CDHZ/W*^21^ or *HINTW*^22,23^ in different samples, *e.g.*, blood, skin or feathers. However, embryo blood and feather sampling is highly invasive and usually fatal. Contrary to this, obtaining an AF sample (a renal filtrate excreted from the embryo during development that contains DNA), is less invasive enabling to collect up to 200 µL on day 9 of incubation without affecting the embryo’s development^15^. Nonetheless, despite such great potential of using AF sample for *in ovo* sexing, scientific reports about DNA detection in AF are still lacking (He et al., 2019; Li et al., 2015).

Therefore, in this work, we develop for the first time an AF-based quantitative PCR (qPCR) for the amplification of ISA Brown chicken female-specific gene from the W chromosome (*HINTW*) as well as for a gene present in the Z chromosome (*DMRT-1*) as a positive control. *HINTW* is a *DMRT-1* gene inhibitor during sexual development. It has several copies (approximately 40) in the W chromosome^24,25^ and is highly conserved in avian species. The use of such conserved genes allows sexual sorting in other chicken lines and bird species due to their shared genetic background^22–24^. To accomplish our objectives, we develop novel primer pairs for the *HINTW* and *DMRT-1* genes using the NCBI-BLAST engine. A comprehensive optimization process is carried out for the qPCR assay, utilizing synthetic DNA and genomic DNA (gDNA) extracted from blood samples collected on incubation day 14. Subsequently, the optimized primers and assay conditions are employed to detect *HINTW* and *DMRT-1* genes in AF-extracted gDNA from samples collected on incubation days 6, 7, 8, and 9. The resulting cycle threshold (Ct) values from the obtained amplification curves and the relation between the Ct values for different genes are analyzed and compared between sexes and sampling days. Moreover, we evaluate the impact of the procedure on embryo development, for each sampling day, by assessing the ratio of dead embryos to the initial number of fertilized eggs incubated and measuring the weight of the yolk and embryo. Using an excretion product of the developing embryo, AF (an understudied sample), we enabled a highly optimized and accurate qPCR assay for determining the sex of chicken embryos early in incubation (as soon as day 6). We also provide with an in-depth study of the possibility of using a relation between *HINTW* and *DMRT-1* genes, for achieving 100% accuracy sexing, with minimal influence to the in-development embryos. Finally, we discuss the possibility of using these results to further improve the current practices seen in *in ovo* sexing DNA-based technologies.

## 2. Materials and methods

### 2.1. Primer design

The primers for *HINTW* (GenBank accession number: NC_052571) and *DMRT-1* genes (GenBank accession number: NC_052572) for chicken (*Gallus Gallus*) were selected using NCBI’s BLAST engine, ensuring specificity for these sequences (Table 1). Using the IDT OligoAnalyzer^TM^ 3.1 (IDT, Leuven, Belgium), the primers’ annealing temperature, stability and self-complementarity were also considered when designing the primers in order to avoid non-specific reactions with the rest of the gDNA or formation of primer-dimers secondary structures. Alignment of the primers with the genes was performed using the MUSCLE software^26^. IDT technologies, USA, produced all the primers and synthetic DNA, the latter in the format of double-stranded DNA sequences (Table 1).

**Table 1.**
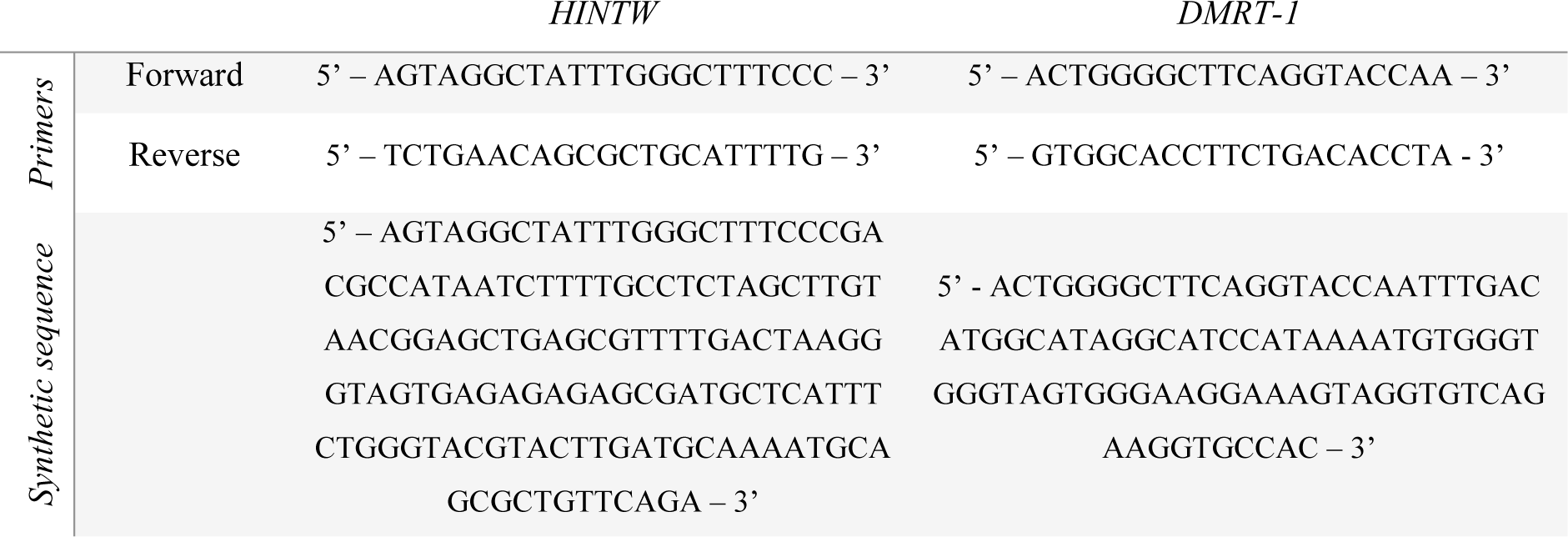
Primers and synthetic DNA used in this paper for the targeted genes.

### 2.2. Materials

The fertilized eggs were from the ISA Brown line and were obtained from a commercial supplier (Vepymo n.v., Belgium). Accu Check® Safe-T-Pro Plus lancing device (Roche AG, Switzerland, Basel) was employed to prick the eggs, from which the AF was extracted using the PRE needle Regular Hub® 32 G (TSK GmbH, Germany, Hanover). gDNA was purified from extracted blood and AF using PureLink™ Genomic DNA Mini Kit (Invitrogen Inc., USA, MS). For the qPCR, the iTaq™ Universal SYBR® Green Supermix (Bio-Rad Laboratories Inc., USA, CA) was used as the master mix while the qPCR was performed using a RotorGene® (Qiagen N.V., Germany, Hilden). Before sequencing, the amplification product was purified using the PureLinkTM PCR purification kit (Invitrogen Inc.). Finally, the amplicons were sequenced using the Rapid Barcoding Kit V14 (Oxford Nanopore Technologies plc, Oxford, UK), for which the reagents were loaded into a SpotOn flow cell (Oxford Nanopore Technologies plc) and read with MinION device (Oxford Nanopore Technologies plc).

### 2.3. Egg handling

Two distinct groups of experiments were carried out: GROUP_1 was used for optimization, accuracy analysis of the developed qPCR and hatchability impact observation, whereas GROUP_2 was used to validate the sexing method in a blind manner thus avoiding biases in the analysis of the results. Importantly, GROUP_1 also comprised a control group of eggs, which remained untouched until day 14 of incubation. Moreover, in GROUP_2, the gDNA extraction from AF and qPCR was performed before visual sexual identification. The parent flocks were respectively 65 and 52 weeks old for these two groups of experiments. The summary of the number of eggs used in each group can be found in Table S1 (Supplementary Material Section S1).

The eggs were numbered using a pencil and stored at 18 °C, 60% relative humidity for 5 (GROUP_1) and 7 (GROUP_2) days before the incubation started. The eggs were then incubated in a Rcom maru max 380 digital incubator (Autolex Co., South Korea) at 37.7 °C, with 55% relative humidity and tilted every hour for 14 days. The incubation was only interrupted on days 6, 7, 8 and 9 for the necessary time to perform the AF sampling by hand (approximately 10 minutes, see Section 2.4 for more details). No egg was sampled for AF more than once. On day 14, the eggs were removed from the incubators, 1) weighed and used for 2) blood drawing from the chorioallantoic vessel before 3) euthanizing the embryos with decapitation. Subsequently, the 4) embryo body and the yolk were separated and weighed to calculate the embryonic development stage. The percentage of dead embryos was calculated by dividing the number of eggs initially present to be sampled on a specific sampling day by the number of dead embryos for the same day upon opening (on day 14). The yolk and embryo weight percentage was obtained by diving those with the egg’s total weight before opening. Finally, the embryos’ corpses were used for 5) visual sexual identification by the feathers’ colors and gonads. The Animal Ethics Committee of the KU Leuven approved all the experiments involving eggs and embryos with project number ECD 134-2016.

### 2.4. Collection and processing of AF and blood samples

AF and blood samples were collected from the chicken embryos at different developmental stages as specified above and further used for DNA analysis. For the AF sampling, the eggs’ handling was done carefully, following the protocol of Weissmann et al., 2013. Briefly, once the eggs were outside the incubators, the air chamber was identified using an egg light candler and a small pencil mark was made 2 mm beneath this chamber, where a small opening (≈ 0.5 mm) was made using Accu-Check^®^ Safe-T-Pro Plus lancing device (Roche AG) pricking needle. The egg was then positioned with an angle of a 45°, keeping the hole at the highest point of the shell surface, and a PRE needle Regular Hub^®^ 32 G (TSK GmbH) with a 4 mm length was used to sample 100 µL of AF per egg.

Blood sampling was performed on day 14, before euthanasia. The chorioallantoic vessel was identified using an egg light candler through the shell and a small window (1 × 1 cm) was opened around it. A total of 200 µL of blood per egg was drawn using a 30 G needle and kept in EDTA sampling tubes.

Both the AF and blood samples were instantly placed on ice during the sampling and stored at −80 °C until further use. gDNA extraction from both type of samples was done using PureLink™ Genomic DNA Mini Kit (Invitrogen Inc.). The DNA concentration and 260/280 ratios were assessed using a NanoDrop 2000 (Thermo Fisher Scientific Inc, USA, MS). The extracted gDNA was directly used in the qPCR protocol (see Section 3.2 and 3.3).

### 2.5. qPCR optimization

qPCR optimization was achieved using a serial dilution of the synthetic DNA (25 – 0.04 ng/µL) followed by a blood-extracted gDNA dilution, while varying: the primer concentration (200 to 500 nM), initial denaturation time (5 to 10 minutes) and annealing/extension temperatures (50 to 70 °C). *DMRT-1* sequence was used as a positive control since it is present in both female and male samples, while a non-template-control (NTC) was performed by replacing the 5 µL of a sample with 5 µL of nuclease-free (NF) water. The qPCR reactions were performed in a total volume of 20 µL with 10 µL of iTaq™ Universal SYBR® Green Supermix (Bio-Rad Laboratories Inc.), 0.7 µL of the forward primer (350 nM), 0.7 µL of the reverse primer (350 nM), 3.6 µL of NF water and 5 µL of DNA sample (in varying concentrations or dilutions as specified above). Thermal cycling conditions, after optimization, were as follows: initial denaturation at 95 °C for 5 minutes, 40 cycles of denaturation (95 °C for 5 seconds) and annealing/extension (60 °C for 30 seconds). The optimized qPCR protocol was subsequently implemented in all the experiments. For melting curve analysis, steps of 1 °C were taken every 5 seconds, between 65 and 95 °C. The qPCR was performed using a RotorGene® (Qiagen N.V.). Data was analysed by studying the Ct values from amplification curves, *i.e.*, the amplification cycle in which the fluorescence signal surpasses 1.

### 2.6. MinION sequencing

The gDNA was purified from AF samples using the PureLink^TM^ PCR purification kit (Invitrogen Inc.) before performing the sequencing with a SpotOn flow cell (Oxford Nanopore Technologies plc) and a MinION device (Oxford Nanopore Technologies plc). The DNA library was prepared using the reagents from the Rapid Barcoding Kit V14 and followed the manufacturers’ protocol (Oxford Nanopore Technologies plc, SQK-RBK 114.24). Purified gDNA samples (10 μL each) were mixed with a unique barcode (1 μL) from the kit and incubated at 30 °C for 2 minutes, followed by incubation at 80 °C for 2 minutes. Barcoded sequences were pooled, washed using AMPure XP Beads (Oxford Nanopore Technologies plc), and resuspended in 15 μL of Elution Buffer. A mixture of the barcoded solution (11 μL) and Rapid Adapter (1 μL) formed the DNA library. The SpotOn flow cell (Oxford Nanopore Technologies plc) was loaded with 1) priming mix, including Flow Cell Flush (1170 μL), Bovine Serum Albumin (5 μL, 50 mg/mL), Flow Cell Tether (30 μL) and 2) with library comprising Sequencing Buffer (37.5 μL), Library Beads (25.5 μL) and the prepared DNA library (12 μL). The SpotOn flow cell (Oxford Nanopore Technologies plc) was inserted into the MinION device (Oxford Nanopore Technologies plc) and the sequencing was performed.

### 2.7. Data analysis

GraphPad Prism version 9 (GraphPad Software, San Diego, CA, USA) was used for all statistical analyses and data visualization. A Shapiro-Wilk test was carried out to check for normality of the data (α = 0.05). While, two-way analysis of variance (ANOVA; α = 0.05), followed by Tukey’s multiple comparisons test, known for its robustness in handling unequal sample sizes was conducted in Sections 3.3, and 3.4, to determine significant differences between group means, in Section 3.6 a one-way ANOVA (α = 0.05) was performed, followed by Tukey’s multiple comparison test.

In Section 3.5, a two sample unpaired Student’s t-test (α = 0.05) was employed to investigate the differences between the means of the two sexes Ct values, allowing us to evaluate whether there were statistically significant differences between them. Additionally, we utilized receiver operating characteristic (ROC) curve analysis to assess the performance of different values in distinguishing between the groups’ sexes. By plotting the sensitivity against 1-specificity, the ROC curve helped determine the optimal cut-off values for maximizing the sexing accuracy of the developed sexing model.

## 3. Results and discussion

### 3.1. qPCR assay optimization with synthetic DNA

The qPCR assay was first optimized to achieve a highly sensitive (≈ 0.04 ng/μL) and specific assay with high efficiency and minimal non-specific amplification. As described in Section 2.5, the optimization involved 16 different testing settings of a series of synthetic DNA dilutions (25 – 0.04 ng/µL), varying primer concentrations (200 – 500 nM), different initial denaturation time (5 – 10 minutes) and annealing/extension temperatures (50 – 70 °C). NTCs were included by replacing the synthetic DNA with water. The qPCR assay composition for both genes (*HINTW* and *DMRT-1*) was selected from the combination providing the highest efficiency (between 95 and 105%) and R^2^ above 0.996 (revealing a distance of 1.66 amplification cycles between each 1:5 dilution step; Bustin et al., 2009). The obtained conditions (Section 2.5) from the optimization were used in whole the qPCR reactions presented in this work.

Figure 1 shows the results of the used optimized assay with a series of synthetic DNA concentrations (*i.e.,* between 25 ng/μL and 0.04 ng/μL in a 1:5 serial dilution) for both *HINTW* (Figure 1A) and *DMRT-1* (Figure 1B) sequences. Five repetitions were carried out for each concentration to demonstrate the assay reliability. The fluorescent threshold (represented with a dotted line) was used to compare the amplification curves by means of Ct values. The obtained calibration curves revealed an efficiency of 102% and an R^2^ of 0.99 for the *HINTW* and an efficiency of 95% and an R^2^ of 0.99 for the *DMRT-1* when using synthetic DNA. Notably, the NTCs showed no amplification (not represented), indicating that the selected primers for both target genes do not form primer dimer or secondary structures. Melting curve analysis was also performed for further comparison with gDNA extracted from blood samples. From the results in Table S2 (Supplementary Material Section S2), it can be seen that the melting peak temperature of the *HINTW* was 82.4 ± 0.3 °C and the *DMRT-1* 80.6 ± 0.1 °C.

**Figure 1.**
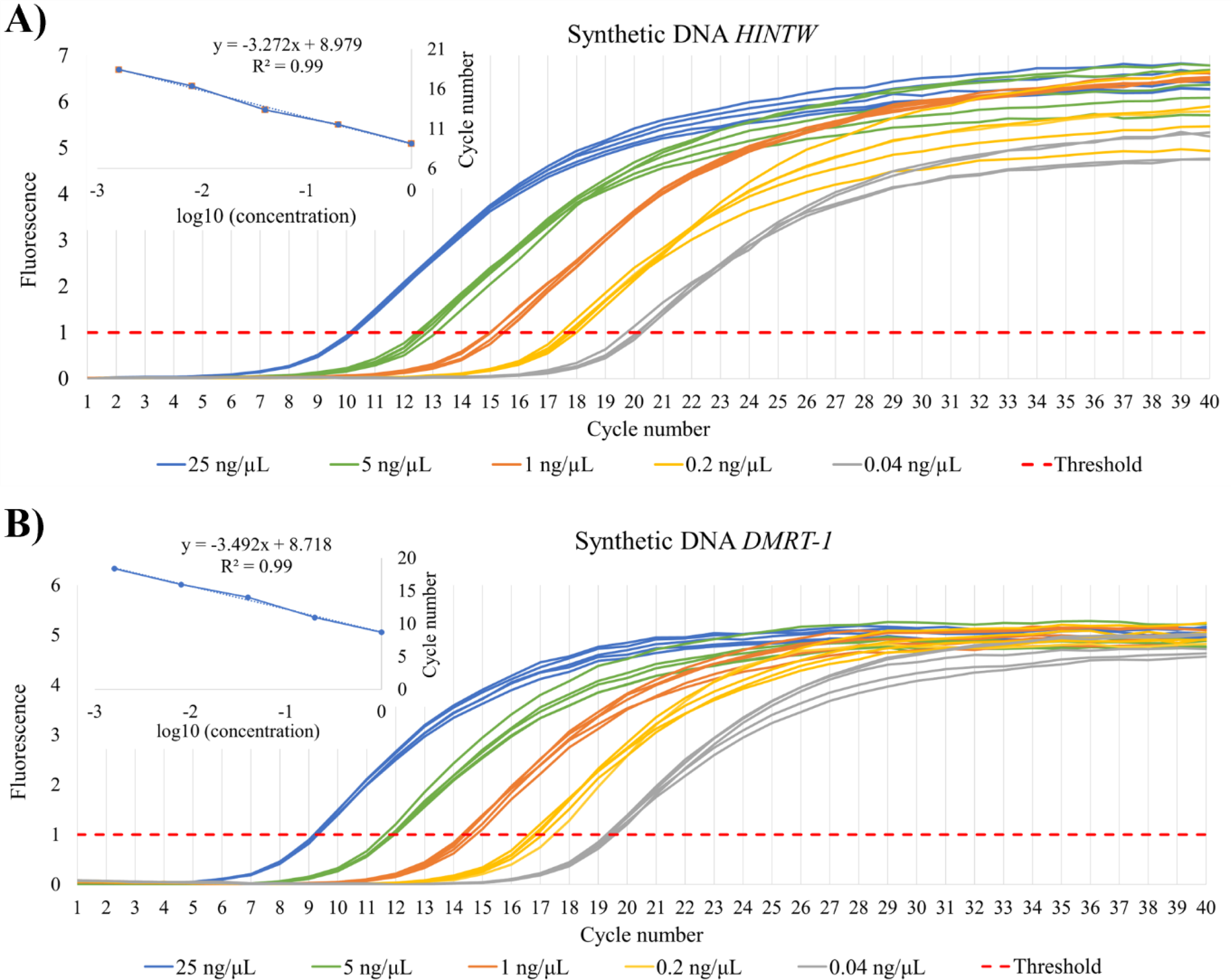
Optimized qPCR reaction performed with serial dilutions of synthetic DNA for A) the HINTW and B) the DMRT-1 genes, revealing R^2^ of 0.99 and efficiency of 102% and 95% for HINTW and DMRT-1, respectively.

### 3.2. qPCR assay with blood-extracted gDNA

In order to test whether the qPCR assay optimized with the synthetic DNA can be used directly for sexing of embryos when the entire genome is present in the AF sample, we tested its performance using gDNA extracted from the blood samples (collected at day 14 of embryonic development from both males and females; Section 2.4). The blood sample was selected here instead of the AF sample because of its abundance and high gDNA concentration from the embryos. The concentration of gDNA extracted from blood was determined using the NanoDrop 2000 (Thermo Fisher Scientific Inc, USA, MS) and subsequently diluted 3 times in a 1:10 ratio. The efficiency and R^2^ of qPCR assay performed with blood-extracted gDNA were calculated for the *DMRT-1* gene (respectively being 95% and 0.99 for females, Figure 2A and 105% and 0.99 for males, Figure 2B), revealing comparable performance to the assay with synthetic DNA (Figure 1). Similar results were observed for the *HINTW* gene in both females (Figure 2C), with an efficiency of 98% and an R^2^ of 0.99 and males (Figure 2D), with an efficiency of 93% and an R^2^ of 0.99. Therefore, we concluded that the obtained qPCR protocol can be directly used with gDNA AF sample without further optimization.

**Figure 2.**
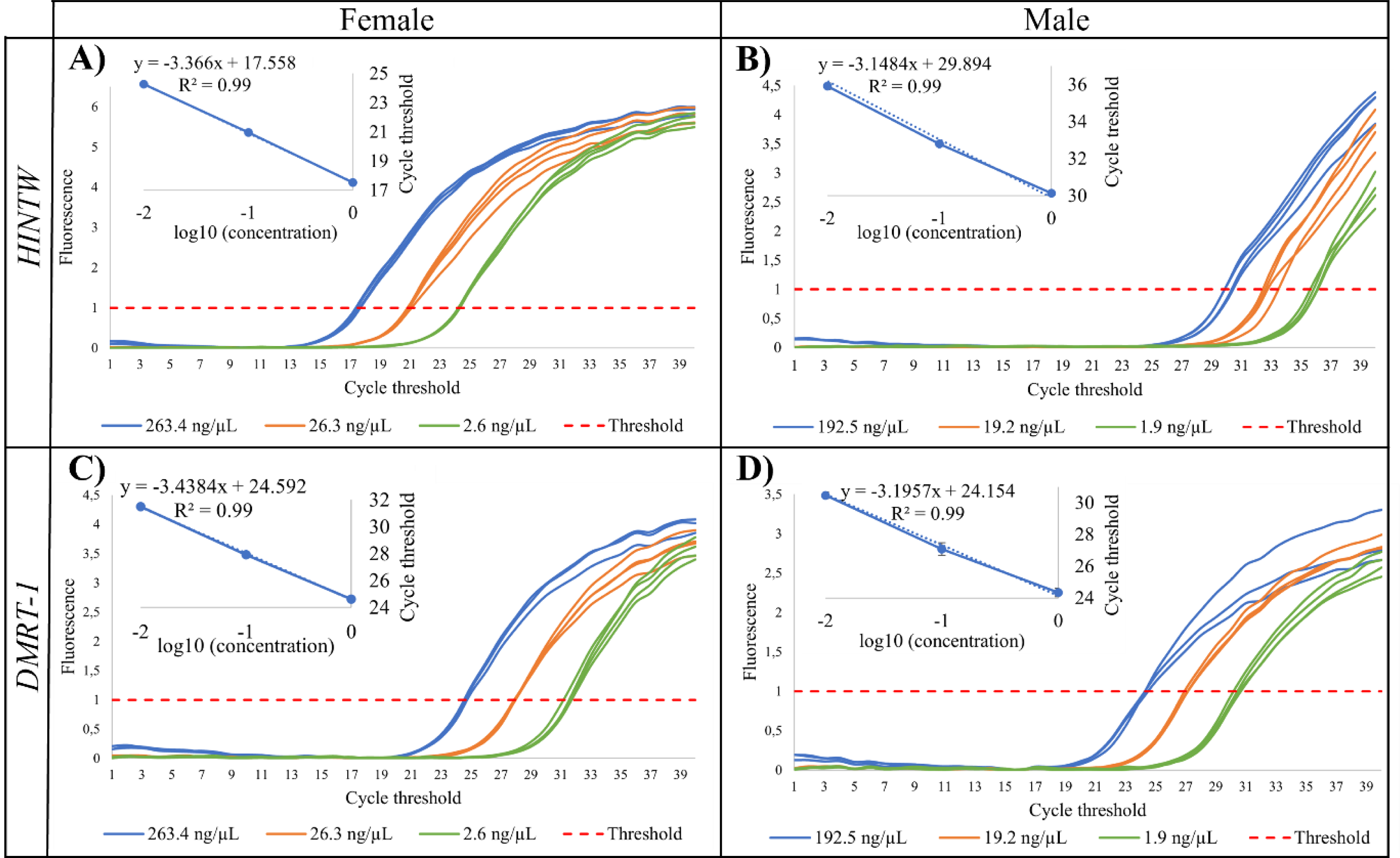
qPCR assay performed with serial dilutions of blood-extracted gDNA from female embryos (shown for DMRT-1 and HINTW in A) and C), respectively) and male embryos (shown for DMRT-1 and HINTW in B) and D), respectively). For females, the R^2^ was 0.99 and efficiency (%) was 98% and 95% for HINTW and DMRT-1, respectively. For males, the R^2^ was 0.99 and efficiency (%) was 93%, and 105% for HINTW and DMRT-1, respectively.

Although not expected, it was also possible to observe some amplification when using the *HINTW* primers with male samples, which had, however, higher Ct values compared to the female samples (Figure 2D). Similar results in blood samples from embryos at day 9 of incubation were described by Cordeiro et al., (2023)^23^, using four distinct chicken lines (Dekalb, Ros, HyLine and Bovan Brown). Thus, we recurred to the melting temperature for more in-depth analysis of these results. From the qPCR assay with synthetic DNA, *HINTW* showed a melting temperature of 82.4 °C (see Section 3.1), which was exactly matching the melting temperature of female *HINTW* gene from blood-extracted gDNA, while it was different compared to 84.2 °C obtained for the male *HINTW* amplification (Table S3 from Supplementary Materials Section S2). Generally, the melting temperature varies among sequences since it relies on their intrinsic characteristics (*e.g.*, sequence length, GC content)^28^. The different melting temperatures between male and female samples show that the sequences amplified in each sex might have differences from each other. To further study the origin of the *HINTW* sequence amplification in male samples, we performed sequencing with gDNA from AF samples, as explained in Section 2.5 and analyzed in Figure S1 of the Supplementary Materials Section S3, showing that the same *HINTW* gene sequence was amplified both in male and female samples, with 94 and 95% identity, respectively. Although *HINTW* gene should not be present at all in the gDNA originating from male chicken embryos, the detected amplification might be explained with the presence of maternal DNA in the egg membranes, as previously described by Strausberger & Ashley (2001)^29^.

### 3.3. qPCR assay with AF-extracted gDNA

To further test whether the established qPCR can be used for AF-based *in ovo* sexing, we collected AF samples at different days of embryonic development (day 6 to day 9) and subsequently divided them into female and male samples based on the visual sexual identification performed at day 14 (for more details, see Section 2.3). In total, 64 eggs were sampled that had an alive and developing embryo at day 14 (Supplementary Materials Section S1). From these 64 eggs, 16 were sampled at day 6 and 7, 13 eggs at day 8 and 19 at day 9. Such divided samples were then used for gDNA extraction and performing qPCR for both *HINTW* and *DMRT-1* gene, of which the amplification curves are depicted in Figure 3. Here, we used only 5 µL of extracted gDNA with a concentration of 7.8 ± 1.5 ng/μL (value retrieved using the NanoDrop from ThermoFisher Inc).

**Figure 3.**
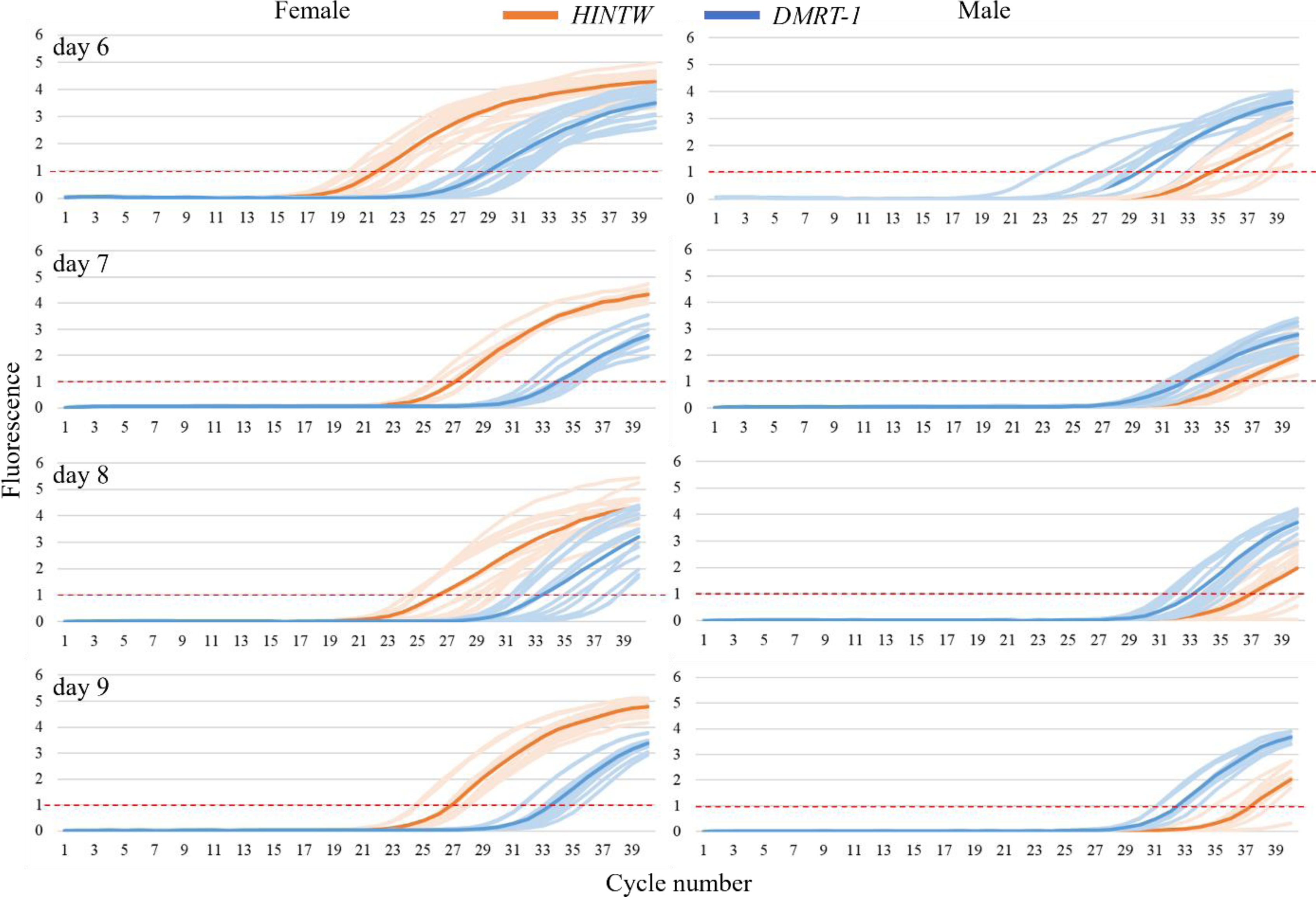
qPCR amplification results from AF-extracted gDNA sampled at different incubation days (day 6, 7, 8 and 9) for both HINTW and DMRT-1 genes. The results show a clear pattern separating male and female chicken embryos. Details about the number of eggs used in these experiments can be found in Table S1 from Supplementary material Section S1.

The average Ct values obtained for the *HINTW* gene from samples collected at different incubation days were plotted for females and males (Figure 4). These results together with the statistical analysis revealed that the Ct values were much higher (> 35 cycles) for the males compared to the females (< 27 cycles), indicating that the concentration of the amplified sequence was considerably lower in the former (further supporting the hypothesis that *HINTW* gene in male samples originates from maternal DNA in the egg membranes). The Shapiro-Wilk test indicated that the Ct values for *HINTW* and *DMRT-1* from each day and sex follow a normal distribution (α = 0.05). Taking into account these results, a Ct value of 31 can be used for *in ovo* sexing (represented with a red dashed line in Figure 4), ensuring a qPCR accuracy of 95%. It is seen that AF samples collected from females on day 6 resulted in a lower Ct value when compared to the female samples from the remaining days, revealing that the gDNA concentrations in this sample are higher on day 6 of incubation compared to days 7, 8 and 9. It is important to note that the proposed Ct value for *in ovo* sexing might have to be re-optimized if the extracted gDNA concentrations in the samples differ. Results related to the cycle thresholds for the *DMRT-1* gene, which has been used as a positive control, can be found in Figure S2 (Supplementary Materials Section S5), showing that the *DMRT-1* Ct values do not significantly differ between males and females for any of the days. These results were expected since the presence of this gene in male and female samples should be approximately the same.

**Figure 4.**
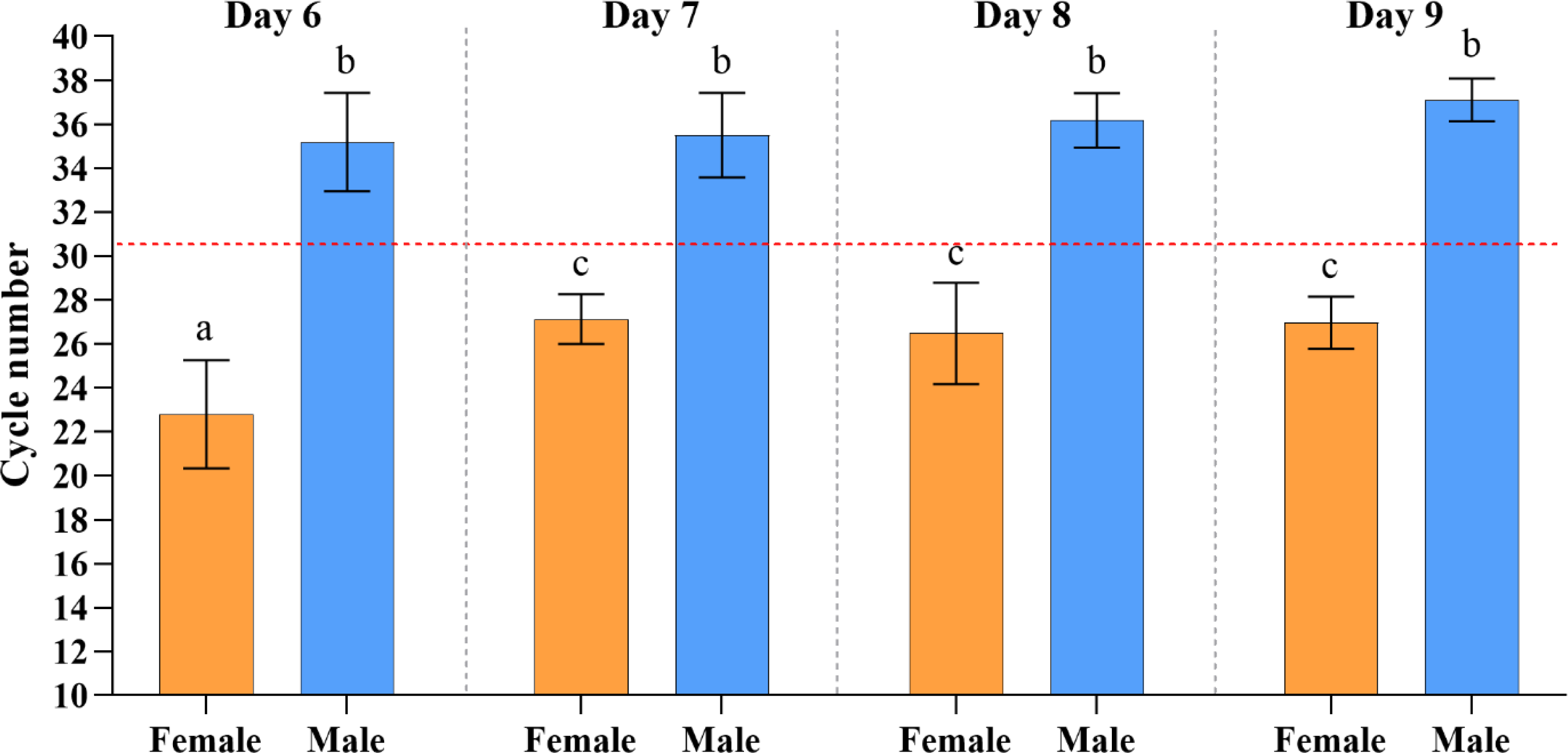
Bar chart represents the mean Ct values obtained from HINTW qPCR amplification curves, for both females and males AF samples collected at different incubation days. The dotted red line represents the proposed Ct value of 31 for in ovo sexing. Error bars represent one standard deviation (n ≥ 6). Statistical analysis involved a two-way ANOVA, followed by Tukey’s multiple comparisons test. While, two-way ANOVA revealed significant differences between sex and sampling days, while the interaction was not significant (α = 0.05). Moreover, Tukey’s tests identified significant differences between 1) females on day 6 compared to days 7, 8 and 9 as well as 2) between females and males for day 6, 7, 8, 9. Different letters above bars indicate significant differences between the means, taking into account multiple comparisons. Details about the number of eggs used in these experiments can be found in Table S1 from Supplementary material Section S1.

Finally, the melting curves were analyzed and the melting peak values (Table S4, Supplementary Materials Section S2) were compared to those previously obtained from synthetic DNA and blood extracted gDNA, revealing no significant differences (Tukey’s test for multiple comparisons).

### 3.4. Relation between *HINTW* and *DMRT-1* amplification thresholds

The developed qPCR is based on two distinct sequences: *HINTW* in the female genome (W-chromosome) and *DMRT-1* in both males and females (Z-chromosome), the latter thus serving as a positive control. In order to further increase the test accuracy and hence not rely only on the Ct value, we used the obtained Ct values for both *HINTW* and *DMRT-1* genes in males and females to calculate Δλ according to Eq. 1 (Figure 5A):

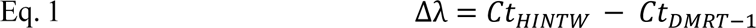

The obtained Δλ values are summarized in Figure 5B and detailed in Table S6 (Supplementary Materials Section S6), revealing that, throughout the sampling days, Δλ is always different between males and females (having a positive or a negative value, respectively). The Shapiro-Wilk test indicated that Δλ follows a normal distribution (α = 0.05) for females and males. A two-way ANOVA analysis was conducted to examine the differences between the mean Δλ for the different days and sexes, revealing significant differences between sexes (α = 0.05). The higher standard deviations obtained for the male samples reflect inherent higher variability during qPCR due to the unspecific amplification of maternal DNA in these samples. The obtained Δλ values also showed that the *HINTW* concentration in females is higher than *DMRT-1*, which is aligned with previous work by Smith et al., (2007)^20^. The same results were obtained with blood-extracted gDNA (Table S7 from Supplementary Materials Section S7), thus confirming that calculating Δλ based on the qPCR Ct values from amplification of *HINTW* and *DRMT-1* genes is a trustworthy method for female identification.

**Figure 5.**
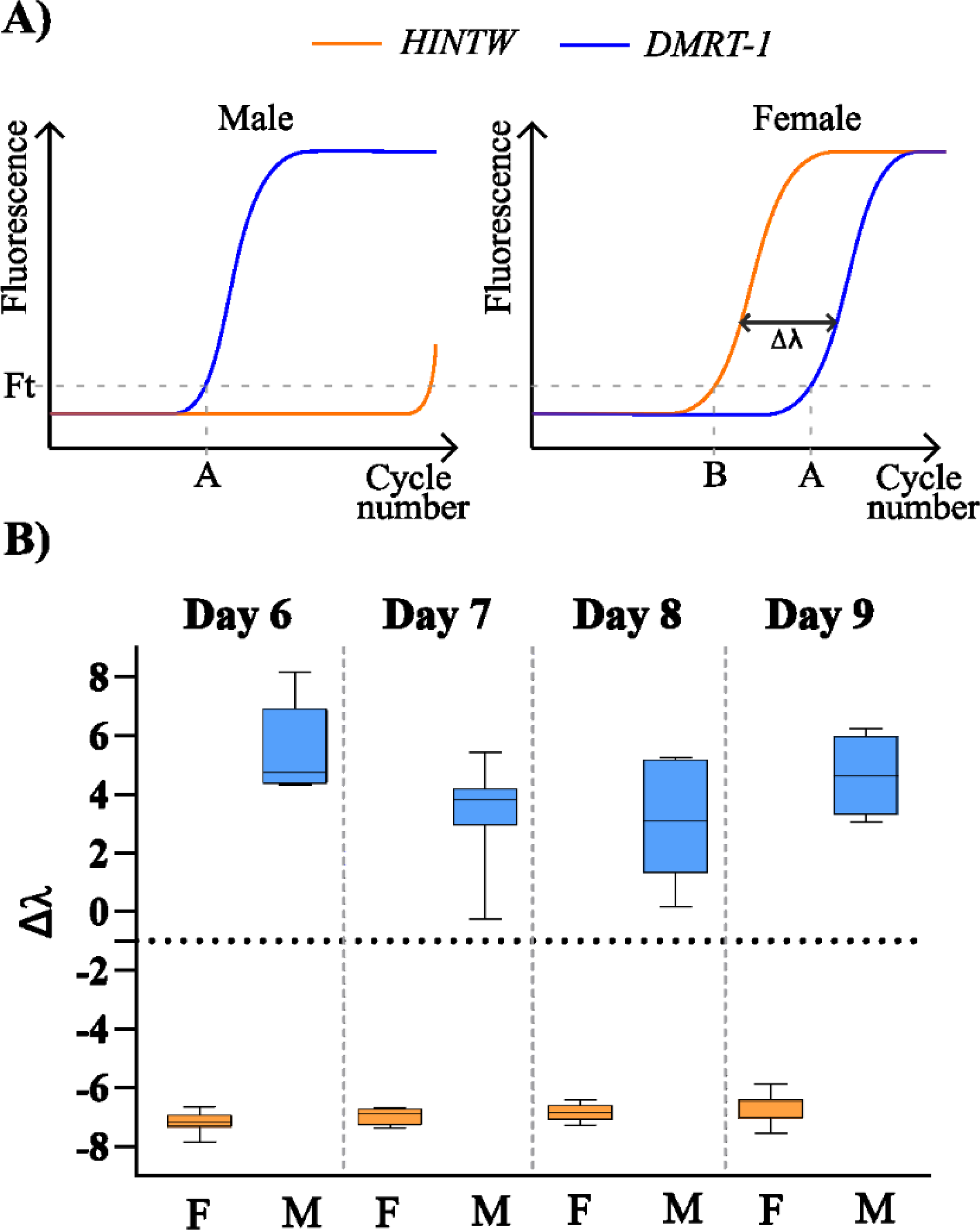
A) Theoretical representations of qPCR amplification curves for HINTW and DMRT-1 genes. The number of samples represented in the graphic are described in the Supplementary materials Section S1. Amplification of DMRT-1 occurs in both sexes. Although HINTW can also be amplified in both genders, in females it has a lower Ct value compared to DMRT- 1, contrary to males where it reveals possible contamination of the eggs. Δλ is represented here as the relation between the genes Ct. B) Values box plot representation of Δλ for different sampling days comparing males with females. Error bars are the 95% confidence interval. Two-way ANOVA revealed significant differences between sexes, while interaction and day was not significant (α = 0.05). Moreover, Tukey’s tests identified significant differences between 1) males on day 6 compared to days 7 and 8 as well as 2) between females and males for day 6, 7, 8, 9, taking into account multiple comparisons. Details about the number of eggs used in these experiments can be found in Table S1 from Supplementary material Section S1.

### 3.5. *In ovo* sexing qPCR assay validation

To validate the accuracy of the established qPCR assay and evaluate its real potential for *in ovo* sexing, we also performed a blind study. 80 eggs were used to sample AF for days 6, 7, 8 and 9 of incubation (20 distinct eggs every day) and the gDNA was extracted from the AF samples (as detailed in Section 2.4) without knowing the embryos’ sex. Shapiro-Wilk test indicated that Δλ, and *HINTW* and *DMRT-1* Ct values for each sex follow a normal distribution (α = 0.05). An area under the curve (AUC) analysis was conducted to quantify the *DMRT-1*, *HINTW* Ct values and Δλ models performance in sexing the embryos without knowing the embryos’ sex, in a blind manner. The AUC analysis allows understanding the discriminatory power of the analysis: with an AUC equal to 1, the model has a strong ability to differentiate between the two sexes, whereas an AUC closer to 0.5 suggests limited discriminatory power. The AUC value is computed by integrating the ROC curve (Figure 6). To validate the findings obtained through the blind analysis, the results were compared with the real outcomes confirmed using gonad inspection on day 14 of incubation (see Section 2.3). The obtained *HINTW* and *DMRT-1* Ct and Δλ values are shown in Figure 6. Moreover, the ROC curve analysis was performed to evaluate the classification model performance, presenting the relationship between the true positive (sensitivity (%)) and false positive (100 – specificity (%)) rates at different Ct for *HINTW* and *DMRT-1* and Δλ value thresholds. Based on these data, it was evident that the *DMRT-1* Ct values could not be used for sexual identification as the values were not significantly different between males and females (unpaired Student’s t test, p > 0.05) with an AUC of 0.57 (Figure 6A). Contrary to this, the average Ct values obtained from *HINTW* gene amplification were significantly different between males and females (unpaired Student’s t test, p < 0.05), with an AUC of 0.9902 showing a sensitivity of 100% and a specificity of 94% for a threshold Ct value of 30.77 (Figure 6B). Lastly, Δλ showed significant differences between female and male samples (unpaired Student’s t test, p < 0.05) and an AUC of 1, providing a sensitivity and a specificity of 100% for a Δλ threshold value of −3 (Figure 6C). The results concluded that the embryo’s sex could accurately be determined by looking at the *HINTW* Ct alone and/or using it together with the Δλ, regardless of the incubation day, since the data analysis was performed by combining samples from days 6, 7, 8 and 9 of incubation (data from the different days is available in the Supplementary Materials Section S8). From this study, we concluded that using *HINTW* could be sufficient for sexing the embryos. However, Δλ can be used to confirm the results further, enabling 100% accuracy in sexing embryos from day 6 to day 9 of incubation.

**Figure 6.**
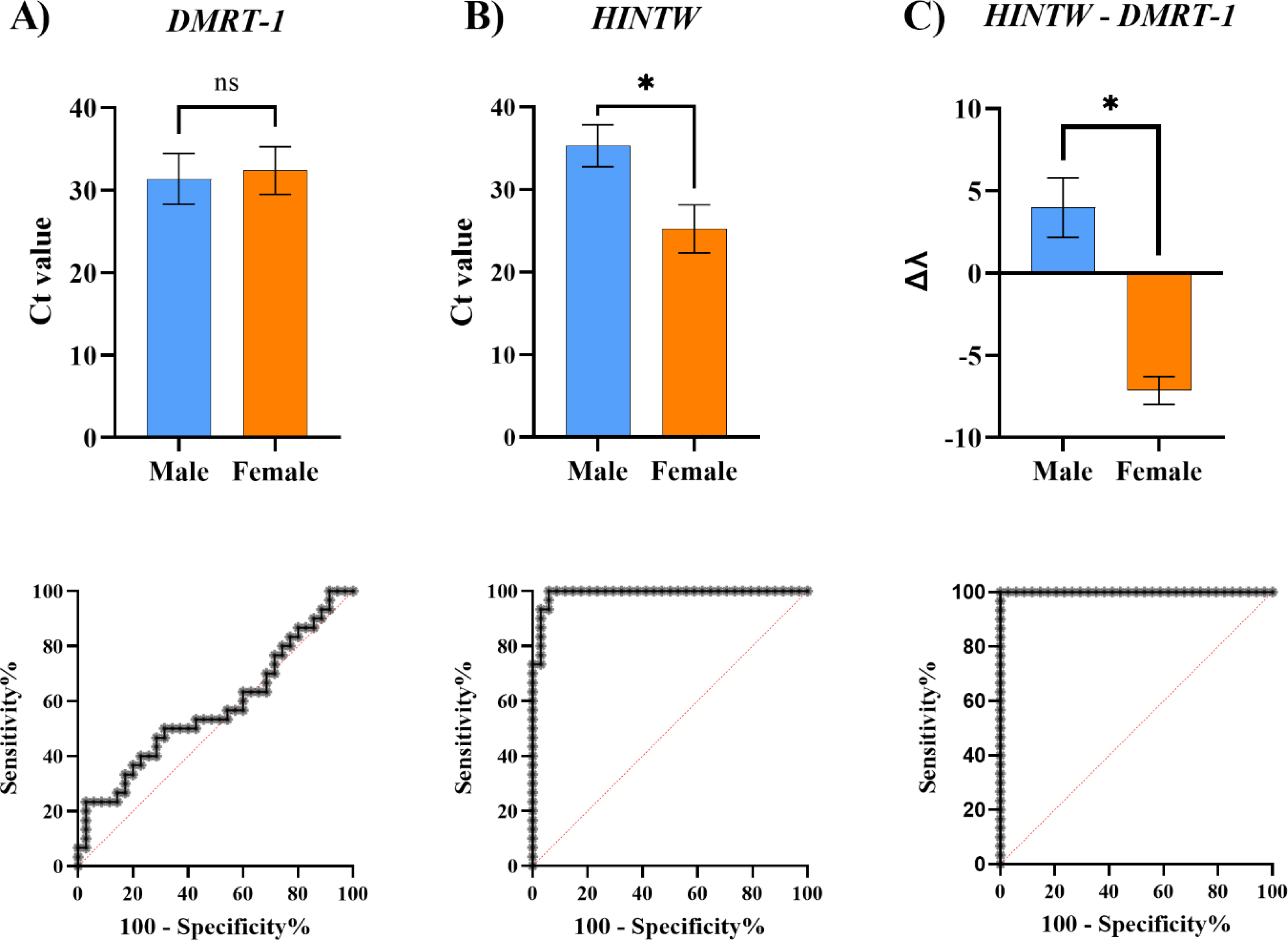
Distribution of Ct for A) DMRT-1 and B) HINTW genes and C) Δλ values obtained in a blind study with AF sampling from eggs at days 6, 7, 8 and 9 of incubation. Error bars represent one standard deviation. The number of samples represented in the graphic are described in the Supplementary materials Section S1. Respective ROC curves are represented beneath. The Ct values of DMRT-1 were not significantly different between males and females (p > 0.05), with an AUC of 0.57. In contrast, the average Ct values of the HINTW gene exhibited significant differences between males and females (p < 0.05), with an AUC of 0.99. A threshold Ct value of 30.77 yielded a sensitivity of 100% and a specificity of 94%. Additionally, Δλ showed significant differences between female and male samples (p < 0.05), with an AUC value of 1. A threshold value of 3 for Δλ provided a sensitivity and specificity of 100%. ns – not significant; * - p < 0.05. Details about the number of eggs used in these experiments can be found in Table S1 from Supplementary material Section S1.

### 3.6. AF sampling and visual sexual sorting

One of the requirements for *in ovo* sexing technologies to be applied in the market is hatchability maintenance and low disturbance of embryo development. In order to evaluate the impact of the AF sampling, we used GROUP_1 to determine the percentage of dead embryos at day 14 of incubation (for details, see Section 2.3) among the eggs used for sampling AF at days 6, 7, 8 and 9 of incubation and a control group, *i.e.*, eggs that remained untouched in the incubator during the 14 incubation days. Moreover, we examined the ratio of embryo or yolk weight over the total egg’s weight as a well-known indicator for appropriate embryo development^30^ for each sampling group. Here, upon opening the eggshell, the embryos and yolks were carefully separated and weighed, as referred to in Section 2.3. Shapiro-Wilk test indicated that embryo weight/total weight of the egg (%) and yolk weight/total weight of the egg (%) ratios for each day follow a normal distribution (α = 0.05).

Figure 7 shows the statistical analysis of the obtained measurements. The eggs for which the AF sampling was performed at day 6, revealed the highest percentage of dead embryos (15.2%; Figure 7A). The lengthier manipulation to extract the AF at this stage might explain this higher percentage, although no clear trend was visible for other sampling days or the control group.

**Figure 7.**
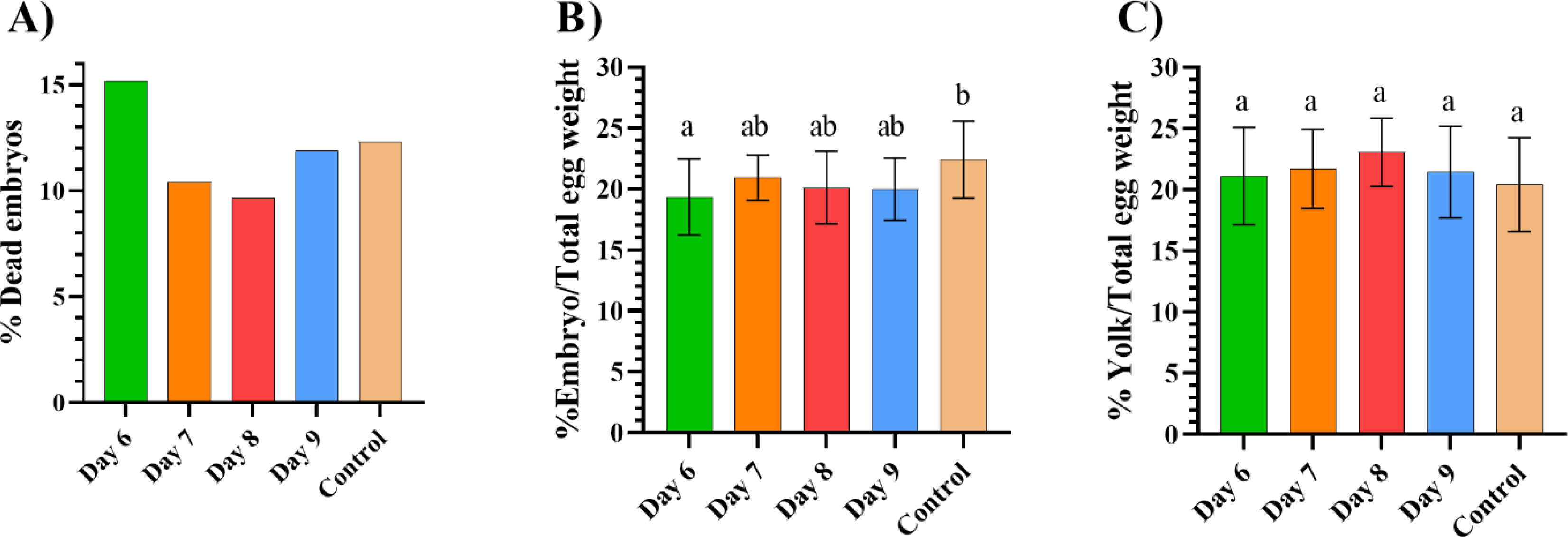
Sampling impact on the sampled eggs compared to a control group. Error bars represent one standard deviation. The control group underwent identical conditions as the sampled eggs, except for uninterrupted incubation until day 14 and the absence of any sampling during the process. The evaluation of the sampling impact was based on A) hatchability (% of dead embryos) and impact on the embryo development by measuring B) embryo weight and C) yolk weight, both relative to the weight of the whole egg. Statistical analysis involved an one-way ANOVA, followed by Tukey’s multiple comparisons test. In B) ANOVA revealed significant differences between groups, α = 0.05), while Tukey’s test revealed significant differences between day 6 and control groups, taking into account multiple comparisons. In C) ANOVA did not reveal significant differences between groups (α = 0.05). Details about the number of eggs used in these experiments can be found in Table S1 from Supplementary material Section S1.

In addition to this, Figure 7B shows that the embryo weight/total weight of the egg (%) was constant among the eggs that were subject to AF sampling at days 7, 8 and 9, and not significantly different from the control group using Tukey’s test (α = 0.05) after one way ANOVA. However, the eggs for which AF sampling was performed at day 6 were significantly different from the control group for this metric (Tukey’s test for multiple comparisons), although this was not the case when looking at the ratio of the yolk weight/total weight of the egg (%) (one-way ANOVA, p > 0.05; Figure 7C). Based on this, we concluded that the AF sampling did not significantly impact embryonic development when compared to the control group of eggs, except for day 6, where the effect on the embryo weight was also minor. In this context, it is important to note that the AF sampling on day 6 has proven to be more challenging compared to the other days, mainly because the AF sample volume on day 6 was limited and variable. From what was observed, it was crucial to wait for some time (≈ 8 minutes) with the egg tilted at 45° for the sample to concentrate at the egg’s top. However, with advancements in user practices, machine automation and even imaging mechanisms such as magnetic resonance imaging, the success rate of AF sampling might significantly increase, especially during the early embryonic development stages, while decreasing the negative effects on hatchability.

## 4. Conclusion

In this work, we developed a reliable and highly accurate method for sexual sorting of female and male eggs between days 6 and 9 of incubation using qPCR assay. The qPCR assay, firstly optimized with synthetic DNA, revealed high efficiencies for amplifying *HINTW* and *DMRT-1* in blood- and AF-extracted gDNA. Using this protocol, the sexual sorting was possible by 1) limiting the number of qPCR cycles to 31 to observe the *HINTW* gene amplification presence only or 2) distinguishing the pattern (represented by Δλ) from the amplification of both *HINTW* and *DMRT-1* genes in males and females. Both approaches revealed remarkable accuracy rates of 95% or 100%, respectively, when a blind study was performed. Although qPCR method is typically regarded in poultry industry as costly, considering the reported accuracy and sensitivity of developed qPCR assay that allow *in ovo* sexing already at day 6, the cost savings can be achieved in the long term by cutting on the energy spending and having higher accuracy. Moreover, additional optimization and novel amplification assays can further reduce the cost, providing even a better prospect for a market application. Lastly, AF sampling on any of the days of incubation (except day 6) did not show significant alterations to embryonic development. However, AF sampling on day 6 of incubation proved to be more challenging than the standard on day 9 due to the variability in AF volume and position, which can be further improved with sampling automation and practices already available at the industrial settings.

## Abbreviations

HINTW: Histidine triad nucleotide binding protein W
DMRT-1: Doublesex and mab-3 related transcription factor 1
qPCR: quantitative Polymerase chain reaction
Ct: Cycle threshold
GMO: Genetic modified organism
AF: Allantoic fluid
ELISA: Enzyme-linked immunosorbent assay
gDNA: genomic DNA
ROC: Receiver operating characteristic
AUC: Area under the curve

## Acknowledgements

This work has received funding from the Flemish environment department and the Research Foundation – Flanders [SB project 1SC7219N and SB project 1S54823N]. Furthermore, gratitude is expressed to Victor Golyaev (KU Leuven, Department of Biosystems, Willem de Croylaan 42, B-3001 Leuven, Belgium), for supporting the sample sequencing.

## Declaration of AI assisted technologies in the writing process

During the preparation of this work the authors used Chat-GPT in order to improve the manuscript readability and succinctness of the Materials and Methods section. After using this tool, the authors reviewed and edited the content as needed and take full responsibility for the content of the publication.

## Notes

### Competing Interest Statement

The authors have declared no competing interest.

