## Supplementary Materials for "Allantoic fluid-based qPCR for early-onset *in ovo* sexing"

### S1 - Number of embryos used in the experiments

*Table S1 – Number of eggs used in the paper for the two groups of experiments. * - Number of eggs used for AF sampling on a given sampling day. ** - Number of eggs with alive embryos used for visual sexual identification and blood sampling. Variation between days in GROUP_1 for AF sampling is related to the presence of unfertile eggs (due to the obtained batch of eggs) at the beginning of the experiment. The final number of eggs used for visual sexual identification and blood drawing on day 14 can be different from the one used for AF sampling due to the death of embryos. A control group (C) was used in GROUP_1, where the eggs remained incubating for 14 days without any interference. GROUP_2 shows that a higher number of eggs was first sampled. However, only 20 eggs per day were included in the study to obtain a homogenous number for each day. NA, not applicable.*

| *Experiment* | *Eggs used for AF sampling** | | | | | *Eggs used on day 14*** | | | | | *Total egg number* |
|---|---|---|---|---|---|---|---|---|---|---|---|
| *Sampling day* | 6 | 7 | 8 | 9 | C | 6 | 7 | 8 | 9 | C | |
| *GROUP_1* | 19 | 18 | 15 | 15 | 22 | 16 | 16 | 13 | 19 | 19 | 89 |
| *GROUP_2* | 25 | 25 | 25 | 25 | NA | 20 | 20 | 20 | 20 | NA | 100 |

### S2 - Melting peak analysis

*Table S2 – Melting peak temperature of the synthetic DNA*

| *Concentration (ng/µL)* | *HINTW (°C)* | *DMRT-1 (°C)* |
|---|---|---|
| *25* | 82.18 ± 0.07 | 80.62 ± 0.08 |
| *5* | 82.28 ± 0.30 | 80.59 ± 0.09 |
| *1* | 82.20 ± 0.34 | 80.57 ± 0.08 |
| *0.2* | 82.49 ± 0.30 | 80.50 ± 0.24 |
| *0.04* | 82.62 ± 0.36 | 80.62 ± 0.16 |
| *Total average* | 82.36 ± 0.34 | 80.58 ± 0.06 |

*Table S3 – Melting peak temperature of the blood-extracted gDNA*

| *Concentration (ng/µL)* | *Female* | | *Concentration (ng/µL)* | *Male* | |
|---|---|---|---|---|---|
| | *HINTW* | *DMRT-1* | | *HINTW* | *DMRT-1* |
| *263.4* | 82.76 ± 0.14 | 80.85 ± 0.67 | *192.5* | 84.25 ± 0.31 | 81.49 ± 0.10 |
| *26.3* | 82.78 ± 0.14 | 81.24 ± 0.03 | *19.2* | 84.05 ± 0.04 | 81.60 ± 0.05 |
| *2.6* | 83.11 ± 0.26 | 81.26 ± 0.04 | *1.9* | 84.39 ± 0.16 | 81.65 ± 0.05 |
| *Total average* | 82.88 ± 0.05 | 81.12 ± 0.30 | | 84.23 ± 0.11 | 81.58 ± 0.02 |

*Table S4 – Melting peak temperature of the AF-extracted gDNA*

| | | *HINTW* | *DMRT-1* |
|---|---|---|---|
| *Day 6* | Female | 82.57 ± 0.54 | 80.55 ± 0.66 |
| | Male | 84.18 ± 0.81 | 81.05 ± 1.36 |
| *Day 7* | Female | 82.34 ± 0.29 | 80.77 ± 0.84 |
| | Male | 83.03 ± 0.60 | 80.96 ± 0.17 |
| *Day 8* | Female | 81.88 ± 0.62 | 80.33 ± 1.73 |
| | Male | 82.92 ± 0.61 | 80.13 ± 0.31 |
| *Day 9* | Female | 82.65 ± 0.11 | 80.95 ± 0.06 |
| | Male | 83.23 ± 0.56 | 80.87 ± 0.03 |
| *All days* | Female | 82.36 ± 0.30 | 80.65 ± 0.23 |
| | Male | 83.34 ± 0.50 | 80.75 ± 0.36 |

### S3 - Sequencing of qPCR amplification products

Sequencing was conducted on qPCR amplicons from the AF-extracted gDNA, using 4 male and 2 female samples, the latter serving as a positive control. The obtained sequences (Sbjct line in Figure S1) were aligned against the complete *Gallus Gallus* genome (Query line in Figure S1) from the NCBI database. Figure S1 displays the alignment results from one of the male (Figure S1 A)) and one of the female (Figure S1 B)) samples, indicating that HINTW gene sequence was amplified both in male and female samples (95% and 94% matching identity, respectively). This suggests that a female-specific gene was amplified in the male samples.

A)

| Score | Expect | Identities | Gaps | Strand |
|---|---|---|---|---|
| 255 bits(138) | 2e-66 | 162/173(94%) | 4/173(2%) | Plus/Minus |

```
Query  4        GTACGTACCCAGAAATGAGCATCGCTTTCTCTTCACTTCCCTTAGTCAAAACGCTCAGCT  63
                |||||||||||||||||||||||||| ||||| |||| |||||||||||||| |||||||
Sbjct  2949908  GTACGTACCCAGAAATGAGCATCGCTCTCTCT-CACTACCCTTAGTCAAAACACTCAGCT  2949850

Query  64       CCGTTACAAGCTAGAGGCAAAAGATTATGGCGTCGGGAAAGCC--AATGGCCTGCTGT-C  120
                ||||||||||||  |||||||||||||||||||||||||||||  ||| |||| |||| |
Sbjct  2949849  CCGTTACAAGCTCAAGGCAAAAGATTATGGCGTCGGGAAAGCCCAAATAGCCTACTGTGC  2949790

Query  121      CTCTTTGCAACATCGAGAGGAAACGAGGCGACGGGACAGAGCCTACTAAATTA  173
                |||||||||||||||||||||||||||||||||||||||||||||||||||||
Sbjct  2949789  CTCTTTGCAACATCGAGAGGAAACGAGGCGACGGGACAGAGCCTACTAAATTA  2949737
```

B)

| Score | Expect | Identities | Gaps | Strand |
|---|---|---|---|---|
| 281 bits(152) | 4e-74 | 169/177(95%) | 1/177(0%) | Plus/Minus |

```
Query  2        CAAGTACGTACCCAGAAATGAGCATCGCTTTCTCTCACTTCCCTTAGTCAAAACGCTCAG  61
                ||||||||||||||||||||||||||||| ||||||||| |||||||||||||| |||||
Sbjct  2949911  CAAGTACGTACCCAGAAATGAGCATCGCTCTCTCTCACTACCCTTAGTCAAAACACTCAG  2949852

Query  62       CTCCGTTACAAGCTAGAGGCAAAAGATTATGGCGT-GGGAAAGCCCAAATAGCCTACTGT  120
                ||||||||||||||  ||||||||||||||||||| ||||||||||||||||||||||||
Sbjct  2949851  CTCCGTTACAAGCTCAAGGCAAAAGATTATGGCGTCGGGAAAGCCCAAATAGCCTACTGT  2949792

Query  121      GCCTCTTTGCAACACCAAGAGGAAACGAGGCGACGGGACAGAGCCTACTAAATTACA  177
                ||||||||||||| | |||||||||||||||||||||||||||||||||||||||||
Sbjct  2949791  GCCTCTTTGCAACATCGAGAGGAAACGAGGCGACGGGACAGAGCCTACTAAATTACA  2949735
```

*Figure S1 – Sequencing of the HINTW qPCR amplicon from the male (A) and female (B) (Query) AF-extracted gDNA samples compared with a fragment of the HINTW gene (Sbjct).*

### S4 - Values of Ct obtained from AF-extracted gDNA

*Table S5 – Values of the Ct obtained from the amplification of gDNA extracted from AF sampled at different incubation days. The normality of the data (Shapiro-Wilk test, p>0.05). Details about the number of eggs used in these experiments can be found in Table S1 from Supplementary material Section S1*

| | *Female* | | *Male* | |
|---|---|---|---|---|
| | *HINTW* | *DMRT-1* | *HINTW* | *DMRT-1* |
| *Day 6* | 22.8 ± 2.5 | 30 ± 2.7 | 35.2 ± 2.2 | 29.7 ± 2.1 |
| *Day 7* | 27.1 ± 1.1 | 34.1 ± 1.2 | 35.5 ± 1.9 | 32.1 ± 2.1 |
| *Day 8* | 26.5 ± 2.3 | 33.3 ± 2.5 | 36.2 ± 1.2 | 33.3 ± 1.5 |
| *Day 9* | 27 ± 1.2 | 33.6 ± 1.1 | 37.1 ± 1.0 | 32.5 ± 1.0 |
| *All days* | 25.3 ± 2.8 | 32.2 ± 2.8 | 35.8 ± 1.8 | 32.1 ± 2.1 |
| *p-value* | 0.1551 | 0.3691 | 0.3665 | 0.123 |

S5 - DMRT-1 threshold cycle obtained from qPCR amplification of AF-extracted gDNA

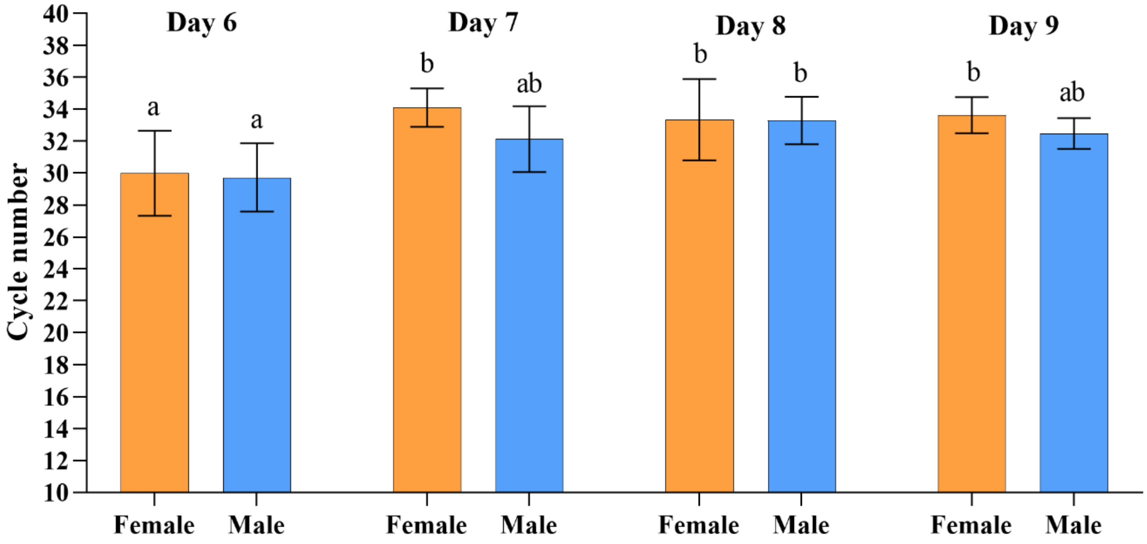

*Figure S2 – Bar chart represents the mean Ct values of DMRT-1, for females and males, for the different incubation days when AF sampling was performed. Error bars represent the standard deviation. The statistical analysis involved a two-way analysis of variance (ANOVA), followed by the Tukey's multiple comparison test. The two-way ANOVA revealed significant differences between sampling days, but not between sex nor for the interaction of sex and day, while interaction was not significant (α = 0.05). The multiple comparison test conducted with the Tukey's test identified a significant difference between 1) females on day 6 and day 7, day 8, day, and 2) males from day 6 and day 8. No significant differences were found for columns with the same letters (α = 0.05). Details about the number of eggs used in these experiments can be found in Table S1 from Supplementary material Section S1.*

### S6 - Δλ values obtained for males and females at different AF sampling days

*Table S6 – Obtained values of Δλ for females and males for the different sampling days (Shapiro-Wilk test, $p > 0.05$). The λ has been calculated based on the Ct values presented in Table S5. Details about the number of eggs used in these experiments can be found in Table S1 from Supplementary material Section S1*

| *AF sampling day (average ± SD)* | *Female* | *Male* |
|---|---|---|
| *6* | -7.19 ± 0.32 | 5.47 ± 1.60 |
| *7* | -6.97 ± 0.29 | 3.38 ± 1.44 |
| *8* | -6.86 ± 0.31 | 3.07 ± 2.07 |
| *9* | -6.66 ± 0.54 | 4.63 ± 1.37 |

### S7 - Δλ value calculations when performing qPCR with blood-extracted gDNA

*Table S7 – Δλ values from the blood-extracted gDNA qPCR amplification for both females and males, calculated based on the Ct values reported in Figure 2*

| *Concentration (ng/µL)* | *Female* | *Concentration (ng/µL)* | *Male* |
|---|---|---|---|
| *263.4* | -7.28 | *192.5* | 7.49 |
| *26.3* | -6.98 | *19.2* | 6.91 |
| *2.6* | -7.15 | *1.9* | 6.35 |
| *Average ± SD* | -6.61 ± 0.65 | *Average ± SD* | 6.77 ± 0.47 |

### S8 – Distribution of Ct for *DMRT-1*, *HINTW* genes and Δλ value in each day

*Table S8 – Ct and Δλ values for female and male for the different sampling days. Details about the number of eggs used in these experiments can be found in Table S1 from Supplementary material Section S1*

| | *Male* | *Female* |
|---|---|---|
| | *Day 6* | |
| *DMRT-1* | 29.72 ± 2.135 | 29.99 ± 2.66 |
| *HINTW* | 35.19 ± 2.24 | 22.79 ± 2.46 |
| $\Delta\lambda$ | 5.34 ± 3.78 | -7.44 ± 1.75 |
| | *Day 7* | |
| *DMRT-1* | 32.13 ± 2.06 | 34.10 ± 1.21 |
| *HINTW* | 35.51 ± 1.93 | 27.13 ± 1.14 |
| $\Delta\lambda$ | 3.38 ± 1.44 | -7.93 ± 2.79 |
| | *Day 8* | |
| *DMRT-1* | 33.29 ± 1.49 | 33.34 ± 2.55 |
| *HINTW* | 36.18 ± 1.23 | 26.48 ± 2.31 |
| $\Delta\lambda$ | 3.70 ± 3.08 | -7.47 ± 3.14 |
| | *Day 9* | |
| *DMRT-1* | 32.48 ± 0.96 | 33.63 ± 1.4 |
| *HINTW* | 37.11 ± 0.97 | 26.97 ± 1.19 |
| $\Delta\lambda$ | 4.63 ± 1.37 | -7.02 ± 0.57 |
